# A model of the cerebellum generates gait adaptations in a reflex-based neuromusculoskeletal model during split-belt walking

**DOI:** 10.1101/2024.12.12.628122

**Authors:** Sophie Fleischmann, Julian Shanbhag, Joerg Miehling, Sandro Wartzack, Carmichael Ong, Bjoern M. Eskofier, Anne D. Koelewijn

## Abstract

**Background:** During split-belt treadmill walking, neurotypical humans exhibit adaptations characterized by a gradual decrease in step length asymmetry (SLA) toward or beyond symmetry, whereas individuals with cerebellar damage do not show these motor adaptations. Neuromusculoskeletal simulations may help to better understand individual aspects of the underlying neural control, but are currently incapable of predicting adaptations to the continuous perturbations imposed by split-belt walking.

**Methods:** We extend a spinal reflex model with a biologically inspired model of the cerebellum, which enables error-based motor adaptation by modulating spinal control parameters in response to mismatches between a continuously updated internal prediction and actual motor outcomes. In this work, the cerebellum modulates only a single spinal control parameter, the timing of swing initiation in each leg, which allows examining its isolated contribution to gait adaptation as all other reflex pathways are held constant. We created 80 s predictive simulations of the model walking on a split-belt treadmill with a 2:1 belt-speed ratio, and compared predicted spatiotemporal parameters and kinematics to the reflex-only model and literature.

**Results:** The reflex-only model could walk on the split-belt treadmill, but showed no step length adaptations. In contrast, the extended model adapted SLA from an initial asymmetric value toward symmetry or beyond, following an exponential time course similar to that observed in experiments. The model could adapt at varying rates and converge to different asymmetry levels. We found that, in simulation, SLA adaptation during split-belt walking is possible without changes in reflex gains, by adapting the timing of swing initiation. The modulation of timing alone also predicted the experimentally observed exponential adaptation in the temporal domain, but only a linear change in the spatial domain, indicating that additional control mechanisms are likely required to reproduce the full spatial adaptation observed in split-belt walking.

**Conclusion:** We propose a computational model of the cerebellum which, when integrated into a spinal reflex model, autonomously drives feedforward gait adaptations during split-belt walking. This advances the current state of predictive simulations and may eventually help to better understand specific adaptation processes. The modular framework can be extended to test different hypotheses about motor control and adaptation during continuous perturbation tasks.

## 1 Introduction

Humans can effectively adjust their gait in response to perturbations or environmental alterations [1]. From a motor control perspective, the observed locomotor responses result from an interplay of different neural control processes within the central nervous system (CNS). Sudden or unexpected perturbations, such as slips, trigger reactive alterations characterized by fast corrective muscle activations in response to sensory feedback [2, 3]. Repeated or continuous exposure to perturbations leads to lasting, error-driven modifications of the locomotor pattern that are associated with motor learning [4, 5].

The most common paradigm for studying locomotor adaptations is split-belt walking [1, 6], where participants walk on a treadmill with the two belts moving at different speeds. Studies usually employ belt speed ratios such as 2:1, 3:1 or even 4:1 [7–14]. Initially, the split-belt perturbation leads to an immediate change in the gait pattern with noticeable spatiotemporal asymmetries, also referred to as feedback- or reactive motor response. However, within several minutes, certain parameters such as the step length or the double support time, gradually adapt toward symmetry [7] or even beyond [14]. These parameters show after-effects once the perturbation is removed, indicating that the new motor plan has been stored, and are considered feedforward-controlled motor adaptations [4, 7, 15]. Feedforward adaptations thus differ from feedback responses in that they emerge gradually and reflect changes in the motor plan that persist even after the perturbation is removed, whereas feedback-driven responses are rapid, transient corrections driven directly by ongoing sensory input to maintain stability [15, 16].

Adaptation studies involving patients with cerebellar damage have highlighted the critical role of the cerebellum in the feedforward adaptation process [15, 17]. Its exact function during motor adaptation is nevertheless a topic of ongoing research. One common hypothesis describes the cerebellum as a forward model whose main functions are sensorimotor prediction and error processing [18, 19]. This forward model has been predominantly studied in the context of effector movement adaptation, particularly in perturbed reaching tasks [20]. The hypothesis suggests that adaptation is driven by a sensorimotor prediction error which emerges from a comparison between a predicted and the actual movement outcome, rather than from discrepancies in target accuracy [21–23]. This error leads to motor corrections and simultaneously updates the internal model from which the prediction is derived [22–24], resulting in the observed feedforward adaptations.

Adaptation during split-belt walking is also considered an error-based process [15, 25], but the error signal driving this adaptation is not as clearly defined as in reaching tasks, where the movement goal is specified by a clear target. Interlimb asymmetry, and particularly step length asymmetry (SLA), has often been proposed as the error signal minimized during adaptation, and is commonly used as a key indicator of the adaptation process [7, 8, 26–28]. However, recent studies have found that individuals can adapt toward positive SLAs [14], challenging the notion of SLA minimization as the underlying error function. Instead, Ishida et al. [29] recently proposed that step velocity asymmetry (SVA) serves as the primary target driving adaptation, which aligns with the observed positive SLAs. They also found that, unlike step length or step time asymmetry, SVA best satisfied the criteria for an optimal control target within the framework of the goal-equivalent manifold, based on their experimental data.

From a biomechanical standpoint, adaptations are characterized by both spatial (foot placement, center of oscillation) and temporal (phasing) adjustments between the limbs [30–34]. Temporal adaptations occur at a faster rate than spatial adaptations [26, 31], and real-time feedback experiments have suggested a hierarchical relationship in which the changes in the temporal influence the spatial domain but not vice versa [32]. Studies focused on muscle activity have further reported adaptations of activation levels [27, 35, 36] and the temporal structure of activation patterns [36]. While the kinematics of the fast and slow leg clearly differ, the individual joint ranges of motion remain largely unchanged throughout the adaptation period [7, 12]. Most studies report the adaptation of interlimb parameters. However, gradual changes during the adaptation period also occur within each limb, such as adjustments in absolute stance time or in the timing of transitions between stance subphases, particularly in the fast leg [7]. For example, we found that the time duration from the fast leg’s heel strike to swing initiation in the same leg increased throughout the adaptation (Figure 1). Swing initiation here refers to the onset of hip and knee flexion and forward leg movement while the foot remains in contact with the ground [37, 38].

**Fig. 1.**
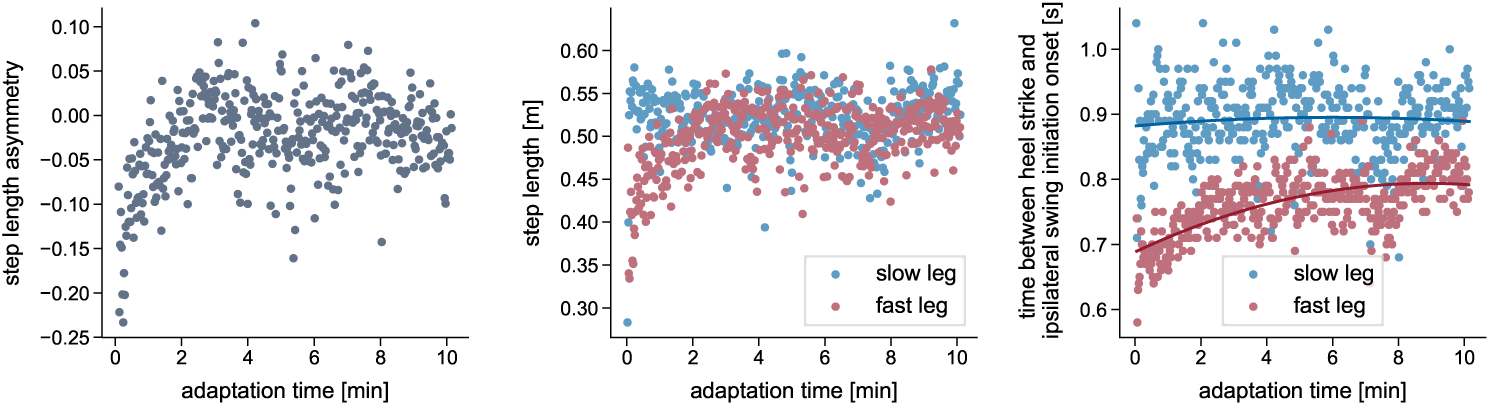
Example of adaptations that occur during split-belt walking. Experimentally recorded data from a representative participant walking on a split-belt treadmill with belt speeds of 1 m s*^−^*^1^ and 0.5 m s*^−^*^1^. The left plot shows the progression of step length asymmetry, calculated stride-by-stride from left and right step lengths, which are shown in the middle plot. The right plot displays the time interval between heel strike and the beginning of swing initiation, which we defined as the time point when the distance between the hip and knee joints began to decrease, indicating the onset of knee flexion and forward leg movement. For details on data recording and computation, refer to S1 File.

The underlying relationships or causal mechanisms between all observed adaptation processes are difficult to determine through experiments, as they often occur in parallel and interact in complex ways. In human experimental studies [32], it is impossible to selectively block individual mechanisms completely, and the ability to probe the function of distinct factors is inherently limited [39–41]. Predictive neuromusculoskeletal simulations provide a key advantage over experimental studies by enabling researchers to isolate and manipulate individual model or control components to assess their contributions and address *what if?* questions [39, 42, 43]. This capability allows to investigate how the change in a single system component, such as a specific reflex or other model control parameter, contributes to motor adaptation. It also enables the systematic examination of the functional consequences when adaptation is selectively absent in specific components, thereby providing deeper insight into the underlying neural control [39, 43]. Simulations can further be used to test different motor control objectives or hypotheses [39, 40].

Gait simulations with reflex-controlled neuromusculoskeletal models have been shown to be robust against different types of mechanical perturbations, such as slips, trips, or drop-down perturbations. However, these perturbations are sudden and discrete and trigger fast, reactive responses [44–46]. In contrast, split-belt tread-mill walking imposes a continuous and sustained perturbation, and induces gradual, cerebellar-driven gait adaptations over time. Most neuromuscular models focus on the spinal control of walking [41, 47, 48], which makes them well-suited for studying reactive responses to brief perturbations, but inadequate for simulating motor learning tasks. Although a few models include supraspinal control structures [49, 50], none incorporate essential physiological components such as the cerebellum, which are required to reproduce and study the gait adaptations observed in split-belt experiments, such as SLA adaptation.

Recently, other areas have begun to apply cerebellum-inspired adaptation mechanisms to dynamic models of locomotion. In robotics, certain aspects of split-belt locomotor adaptations have been qualitatively reproduced using torque-controlled robots [51, 52]. However, it is challenging to transfer these approaches to neuromusculoskeletal models. Furthermore, the insight into human motor adaptation is limited due to differences in system dynamics and control. The same applies to the work by Seethapathi [6], in which changes in SLA were successfully predicted by adding a reinforcement learner onto a biped walker. The reinforcement learner acted as a supraspinal controller and adjusted nominal values of the biped’s controller to minimize energy. But since step length was one of the walker’s two control variables, and a nominal value of step length was also used as input to the controller and adjusted by the cerebellum, the simple model can neither explain physiological reasons for adaptation nor the interaction between the CNS, multi-segment body dynamics, and the overactuated musculoskeletal system during the adaptation process. There is one simulation study focusing on gait adaptations in split-belt walking that uses a neuro-musculoskeletal model [53]. Based on the experimental finding that the gastrocnemius muscle’s H-reflex is modulated during adaptation [54], the authors showed with their simulations that an adaptation of the model’s spinal reflex gains can directly lead to a change in SLA [53]. However, the changes in reflexes in the simulation, although motivated by their previous findings, were externally prescribed as an exponential function, since the model lacked an internal supraspinal control structure that could independently compute them based on the movement.

In this study, we extend a neuromusculoskeletal reflex model with a supraspinal control layer representing the cerebellum, which autonomously drives feedforward gait adaptations during split-belt walking. Changes in control parameters are not prescribed but emerge from the adaptive behavior of the cerebellar model. In our work, we adopt the theory that cerebellar locomotor adaptations are an error-driven process [15, 25]. Our cerebellum model is designed to reflect key physiological functions of the human cerebellum: it predicts the outcome of motor commands, compares the prediction to actual sensory feedback, and uses the resulting prediction error to both adjust outgoing motor signals and update its internal prediction [18, 19, 22, 23]. The spinal control system involves a large number of parameters that interact in complex, nonlinear ways, making it difficult to identify which ones are adapted by supraspinal structures like the cerebellum. Thus, even though our computational cerebellum model is capable of adapting any spinal control parameter, this work focuses on a single adaptation mechanism, the timing of swing initiation in each leg. By restricting adaptation to timing alone and holding reflex parameters constant, we can test whether reflex changes are essential for adaptation or whether adaptation can occur without them, and at the same time isolate the functional role of timing adaptations to clarify their contribution to gait adaptations. We generate predictive simulations using single shooting methods to optimize the asymmetric muscle reflex parameters of a gait controller and the internal cerebellar parameters for split-belt walking at a 2:1 belt speed ratio. We test the effectiveness of the cerebellum model in predicting locomotor adaptations by comparing the SLA at the beginning and end of the simulation to that of a reflex-based model without the cerebellum. The extent to which features of locomotor adaptation are predicted is further assessed by comparing predicted gait parameters to literature. We also analyze changes in kinematics and muscle activations, and investigate how much the adaptations affect the spatial and temporal domains. Eventually, we use our model to examine the effects of the update rates of the cerebellar internal model and motor commands on step length adaptation and on the SLA level at the end of adaptation.

The integration of the cerebellum model led to SLA adaptations in the reflex model similar to those reported in experiments. We show that, in simulation, SLA adaptation during the split-belt walking period is possible without reflex gain changes, though only by adapting the timing of gait phases. However, our results also suggest that the adaptation of additional control mechanisms, like reflexes, may be necessary to fully reproduce the exponential changes observed in the spatial domain of human split-belt adaptation.

## 2 Materials and methods

All simulations are conducted in SCONE [43] (version 2.3.1), an optimization and control framework for predictive simulations, and with Hyfydy [55] as the dynamics engine. Our extended model has a hierarchical control structure consisting of a spinal and supraspinal level, which interact with the musculoskeletal system. The spinal layer computes neural muscle excitation signals based on delayed sensory feedback. These excitations are then translated into muscle forces in the musculoskeletal layer using virtual muscle representations. The supraspinal level, representing the cerebellum, continuously adjusts the spinal control based on an error signal while simultaneously updating its internal model. An overview of the approach is given in Figure 2. The free controller parameters are found by minimizing a multi-objective cost function representing CNS movement goals, but this cost function is distinct from the cerebellar error-based adaptation process. In the following sections, the different layers are described in more detail together with the treadmill objective function and the optimization protocol.

**Fig. 2.**
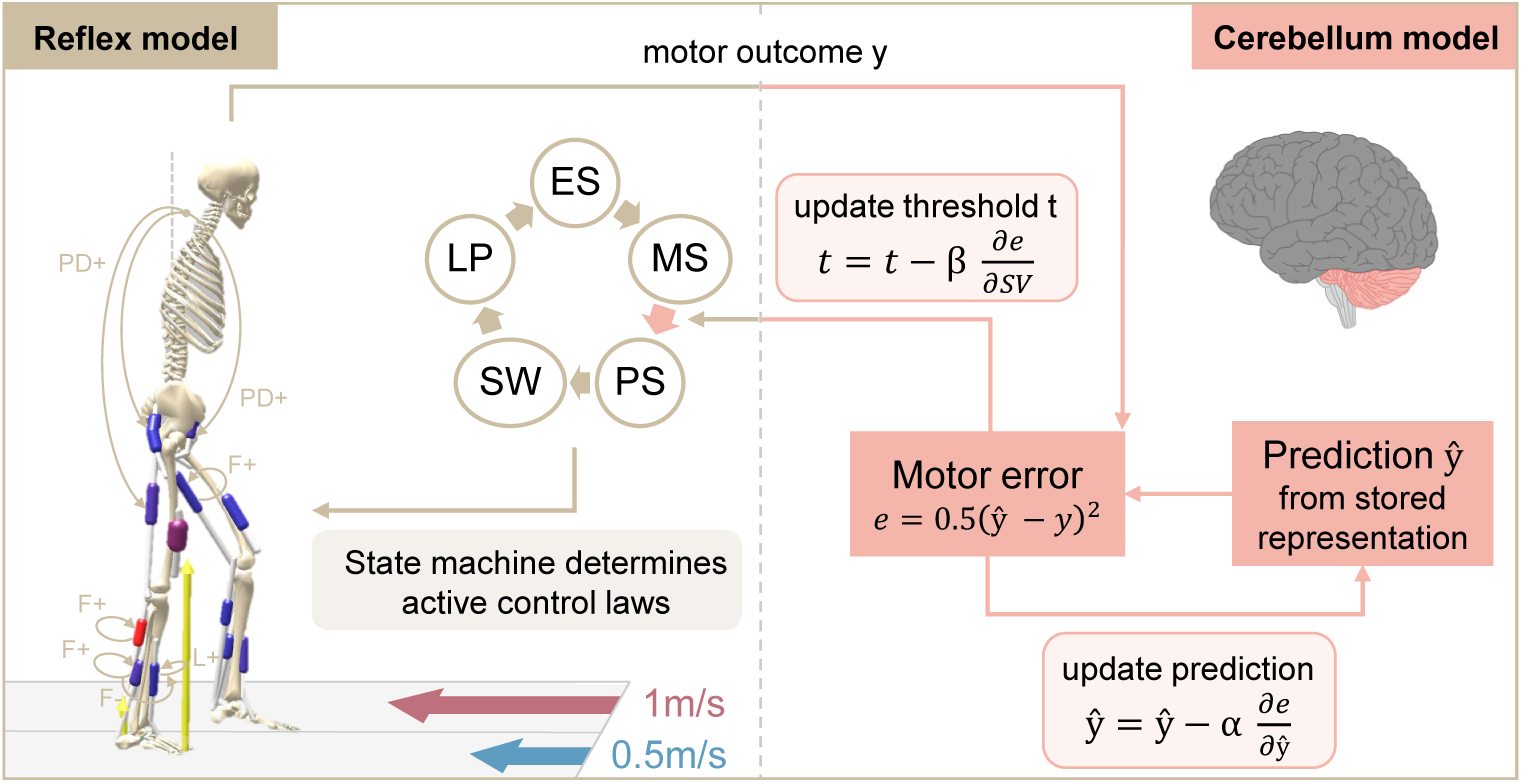
Overview of the spinal and supraspinal control interaction of the extended reflex model. The reflex model has a spinal gait controller based on [56]. A state machine selects the active reflexes based on the current model state, which can be early stance (ES), mid-stance (MS), pre-swing (PS), swing (SW), or landing preparation (LP). The cerebellum compares the motor output *y* (step velocity asymmetry in our model) to a predicted output *y*^, which is generated from an internal representation. Both spinal control and internal prediction are updated based on the partial derivatives of the error with respect to the step velocities SV and to *y*^, at rates *β* and *α*, respectively. In this study, the modulated spinal parameter is the beginning of the pre-swing phase.

### 2.1 Musculoskeletal and treadmill model

The two-dimensional sagittal musculoskeletal model is based on the work from Delp [57] with updates from Rajagopal [58]. It is distributed through SCONE and includes a pelvis-trunk segment and two legs, each composed of thigh, shank, and foot. The body segments are connected via six pin joints representing the hip, knee, and ankle joints. A 3-degree of freedom joint between the pelvis and ground allows for translation in the anterior-posterior and vertical directions and for pelvis tilt, resulting in a total of 9 degrees of freedom for the entire model. The lumbar joint is locked at 8.5*^◦^*of flexion. Each leg is actuated by seven Hill-type muscle-tendon units [59] that represent the major limb muscle groups: hamstrings (HAM), gluteus maximus (GLU), iliopsoas (IL), vasti (VAS), gastrocnemius (GAS), soleus (SOL), and tibialis anterior (TA).

The treadmill model consists of two belts atop the ground. Each belt is modeled as box-shaped contact geometry that could move in the anterior-posterior direction at a predefined speed. Ground reaction forces between the feet and the treadmill belts are computed using the nonlinear Hunt-Crossley contact model [60] and two contact spheres per foot, one at the heel and one at the toes.

### 2.2 Spinal reflex control

The spinal control is based on the reflex-based gait controller from Geyer and Herr [56] (Figure 3). Spinal reflexes translate sensory information into muscle excitation signals via *α*-motorneurons [56, 61]. Our reflex loops incorporate both muscle force and muscle length information, representing the sensory feedback from the Golgi tendon organs and muscle spindles [61]. The trunk is stabilized with proportional-derivative (PD) control as proposed in [41, 48, 61]. In addition, there are certain constant feedforward muscle excitation signals [56]. The control loops are subject to neural delays that range from 5 ms to 20 ms depending on the distance the signal must travel along the nerves between the muscle and the spinal cord. A mathematical description of the spinal control laws is provided in S4 File.

**Fig. 3.**
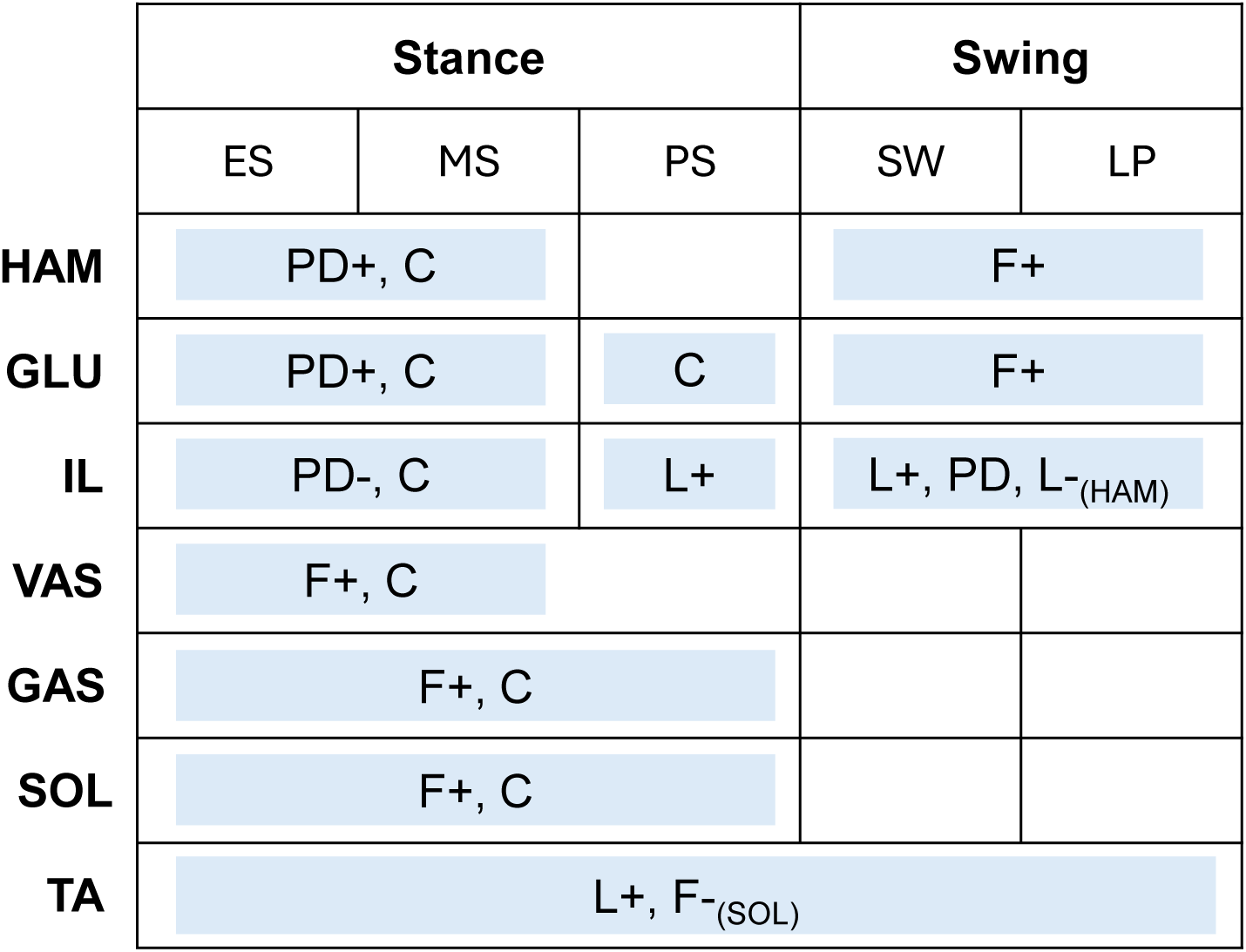
The spinal controller computes muscle excitations from different control laws, which are activated based on a state machine. The state machine determines whether the model is in early stance (ES), mid-stance (MS), pre-swing (PS), swing (SW), or landing preparation (LP). Low-level control laws include muscle force feedback (F), muscle length feedback (L), proportional-derivative control based on the pelvis tilt (PD), or a constant signal (C). The (+) and (-) indicate positive and negative feedback. The subscript means that the signal is received from the respective muscle, if there is no subscript the feedback law acts on the same muscle that it is based on.

Reflexes are generally phase-dependent, meaning they are inhibited or activated during certain periods of the gait cycle [62]. This phase dependency is realized by a state machine that, based on the model’s current state, activates selected spinal reflexes. The possible states for each leg correspond to subphases of the gait cycle: early stance (ES), mid-stance (MS), pre-swing (PS), swing (SW), and landing preparation (LP) [41]. The transition between states is based on established thresholds [48, 63, 64] with one modification that concerns the switch from mid-stance to pre-swing. In simulations of normal walking, this transition, which marks the start of swing initiation, occurs at the moment of contralateral heel strike [63]. However, we found this trigger to be insufficient in simulating split-belt walking, as initiating swing strictly at contralateral heel strike prevents variations in double support duration and limits the model’s ability to capture changes in interlimb phasing and timing. Therefore, we introduced an alternative swing-initiation threshold *t*, defined as a certain delay after the ipsilateral heel strike, as the trigger for the spinal controller to enter pre-swing state. In our work, *t* thus represents the duration between a leg’s heel strike and the start of swing initiation and is a free spinal control parameter included in the optimization.

### 2.3 Cerebellum model

In this study, we adopted the theory of sensory prediction errors driving adaptation and applied it to locomotion. We developed a cerebellar model that maintains an internal prediction, which is updated using prediction errors, and adjusts locomotor patterns based on the error signal. Locomotor adaptation in our framework is thus driven by minimizing asymmetry-related errors rather than optimizing energy expenditure. In this work, we used SVA as the error signal [29]. The cerebellum computes the sensorimotor error *e* as the squared difference between a predicted and the actual observed SVA:

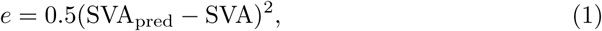

where

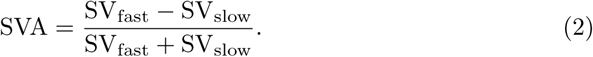

During normal walking, this error signal is zero and no motor corrections are made. At the start of the split-belt perturbation, the error increases, leading to subsequent movement corrections. SV_fast_ and SV_slow_ denote the step velocities of the legs on the fast and the slow belt, respectively. Step velocity is computed as the ratio between step length *l* and step time *d* of the respective leg [29]:

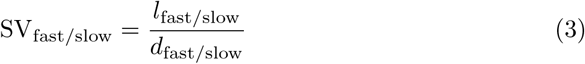

The step lengths are computed as the anterior-posterior distance between both feet at the respective heel strikes (HS) [7], where the calcaneus center of mass (COM) is used to calculate the foot position. Step time for each limb is defined as the time interval between the heel strike of the contralateral limb and the heel strike of the respective limb (e.g., for the slow leg, from fast HS to slow HS) [29]. The observed SVA is updated after every HS.

The prediction [65], as well as the control signals, are updated based on the error via gradient descent shortly after every HS. For the prediction, this corresponds to an update in the direction of the negative gradient of the error with respect to its current value:

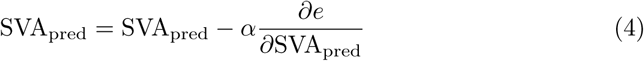

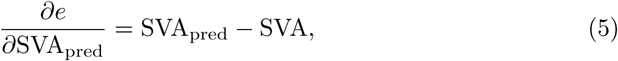

where *α* is the cerebellar learning rate. The initial value of SVA_pred_ is set to zero, corresponding to symmetric walking during baseline tied-belt conditions where no adaptations happen.

The motor control signals are updated in a similar gradient descent manner. In our study, we adapt only timing-related spinal control signals, specifically the start of swing initiation. Accordingly, our cerebellum model adjusts the threshold *t* of each leg, which controls the timing of the leg’s transition from mid-stance to pre-swing. The magnitude of each update is dependent on the gradient of the error with respect to the step velocity of the corresponding limb, scaled by the adaptation rate *β*. The left and right threshold parameters *t*_slow*/*fast_ of the state machine are individually apdated, but based on a single adaptation rate *β* that is shared across both legs:

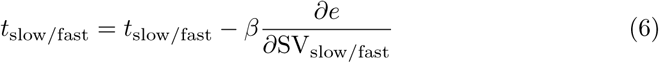

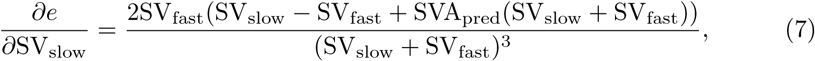

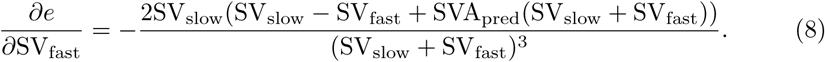

The cerebellar learning rate *α* and the adaptation rate *β* are free controller parameters that are optimized. Since the swing initiation thresholds *t*_slow*/*fast_ are part of the spinal controller, their initial values are also found via optimization. Unlike the approach in [51], our method enables the individual adaptation of both limbs using a single adaptation rate and a single prediction.

### 2.4 Objective function for treadmill walking

All commonly used objective functions are designed for overground walking, where the ground is stationary and the body moves. These functions cannot be directly applied to treadmill walking, where the COM remains relatively stationary while the treadmill belts move beneath the body. We therefore designed a tiered objective function *J*, specifically tailored for treadmill walking, which is sought to be minimized:

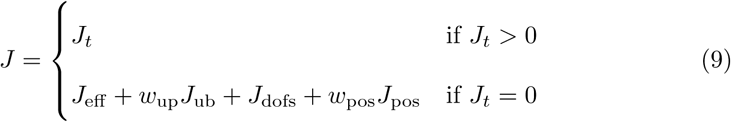

All weight factors *w_j_* are shown in Table 1. The primary objective *J_t_* is used to find an initial solution for the model to maintain its position on the treadmill throughout the desired simulation period *t*_des_ without falling, being dragged backward, or moving forward excessively:

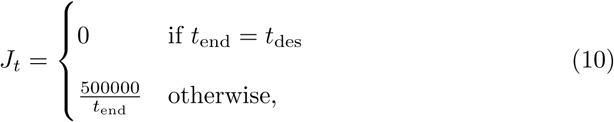

**Table 1.** Weight factors of the individual objective terms.

| $w_{\text{ub}}$ | $w_{\text{pos}}$ | $w_{\text{met}}$ | $w_{\text{act}}$ | $w_{\text{ankle}}$ | $w_{\text{knee}}$ |
| --- | --- | --- | --- | --- | --- |
| 0.05 | 100 | 0.001 | 20 | 0.1 | 0.01 |

The weight factors were selected based on a combination of existing recommendations through SCONE and iterative empirical adjustments necessary to achieve a stable and physiological gait pattern under treadmill and split-belt conditions. Since all objective terms are in different units, a higher weight is not necessarily an indication of higher importance of the respective term.

where *t*_end_ is the total integration time, which continues until the desired simulation time is reached or early stopping is triggered. This happens when the model’s vertical COM drops below 85 % of its initial position (indicating that the model has fallen) or when the anterior-posterior pelvis COM deviates more than 10 m from its initial position.

Once this base condition is met (*J_t_* = 0), a different set of objective terms is evaluated. It consists of several objectives and physiological constraints relevant to human gait in general, and one specifically for treadmill walking, which ensures that the model’s COM remains stationary in the global coordinate system. The objective function is designed to represent the overall goal of walking, assuming that the CNS selects gait patterns that are energetically efficient [66] while simultaneously satisfying other relevant criteria such as stability, or task-specific demands [41, 48, 67]. Specifically, *J*_eff_ minimizes the metabolic rate *E* as defined in [68] and the cubed muscle activations a of the model:

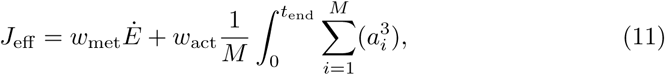

where *M* is the number of muscles and *w*_met_ and *w*_act_ are weighting factors. The second term, *J*_ub_ penalizes excessive upper body and head motion [69], by applying a penalty of 100 whenever the absolute pelvis tilt velocity exceeds 10 ◦ s*^−^*^1^. *J*_dofs_ prevents hyperflexion and overextension of the ankle joints by penalizing the respective joint angle *α* exceeding a range [*α*_min_ *α*_max_], which was [*−*60*^◦^*, 60*^◦^*] in our case [70, 71]. Additionally, non-physiological knee motions are prevented by minimizing the knee limit torque *T* caused by the virtual passive knee ligaments [48].

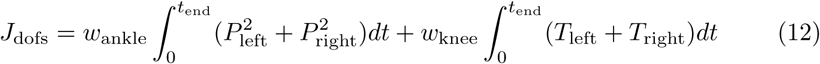

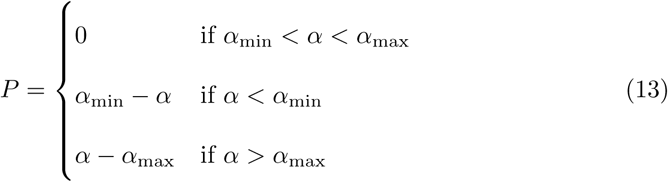

Finally, *J*_pos_ is included in the objective to keep the model in a stationary position, since treadmills have spatial constraints that limit forward or backward movement:

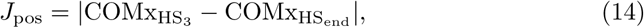

where 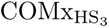 and 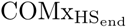 are the model’s anterior-posterior COM position at the third HS and last HS of the same leg, respectively. We used the third instead of the first HS in the objective to allow the model to stabilize first [72].

### 2.5 Generation of split-belt simulations

Our spinal controller was parameterized by 64 free parameters (62 reflex controller parameters (see S4 File) and the left and right threshold parameters *t* from the state machine). The cerebellum model had two additional free parameters, the learning rate *α* and adaptation rate *β*, resulting in 66 free parameters for the extended model.

The covariance matrix adaptation evolution strategy (CMA-ES) [73] was used to optimize these variables. Similar to other studies, we set the CMA-ES parameters to have a population size *λ* of 16 and an initial step size *σ* of 1 [61]. The optimizations terminated either after a maximum number of generations [41] or when the fitness no longer improved, which was the case if the objective function averaged over a window of 500 generations decreased less than 1*e−*4 [70, 74]. The initial generalized coordinates and velocities of the model were set, but not optimized. Initial muscle activations and corresponding muscle states were computed automatically during SCONE’s internal initialization. In this process, the first muscle activations were determined by applying the gait controller’s control laws to muscle-tendon lengths and forces obtained through equilibration of the muscle-tendon dynamics based on default minimal activations. The corresponding initial muscle states were then determined by re-equilibrating based on these newly computed muscle activations.

To find an initial guess for the muscle reflex parameters of the spinal controller for split-belt walking with a 2:1 belt speed ratio, we generated multiple predictive simulations, starting with both belts running with 1 m s*^−^*^1^ and gradually reducing the right belt speed to 0.5 m s*^−^*^1^ in increments of 0.1 m s*^−^*^1^. For each speed ratio, the best result from the previous optimization was used as the initial guess for the next, enabling the model to maintain stable walking while increasing belt speed asymmetry. However, when the 2:1 belt speed ratio was reached, the resulting gait pattern did not exhibit the same level of asymmetry observed in humans at the start of the split-belt perturbation. This discrepancy likely results from the noisy, non-convex optimization landscape [41], in which local optima can prevent the optimizer from reaching more distant regions of the parameter space that would result in more asymmetric gait patterns [75]. To better approximate the initial behavior seen in human split-belt walking, we thus manually readjusted selected reflex values, specifically the GAS and SOL force reflex gains during the stance phase, as well as the IL length reflex, because these parameters have been shown to influence step length [61]. We reran the optimization with belt speeds set to the 2:1 ratio and the adjusted initial guess, and the resulting optimized values were used as the initial guess for all subsequent simulations of the split-belt adaptation period.

Eventually, we generated separate 80 s simulations for the reflex-only and the reflex model extended by the cerebellum. To simulate the adaptation phase, the belt-speed ratio of all simulations was set to a 2:1 belt-speed ratio, with the left belt running at 1 m s*^−^*^1^ and the right belt at 0.5 m s*^−^*^1^. These speeds were chosen based on their previous use in experimental studies. In human experiments, the adaptation phase typically lasts around five to ten minutes, but due to the high computational cost of predictive simulations, we limited the maximum duration of our simulations to 80 s. Given the stochastic nature of CMA-ES and the highly nonlinear and non-convex search space [41], we ran ten parallel optimizations with different random seeds for both configurations, using the same previously determined initial guess and a maximum of 1500 generations. The first gait cycle of the 80 s simulations was excluded from the analysis [48, 72].

Our cerebellum model is parameterized by the learning rate *α* and adaptation rate *β*, where *α* defines the rate at which the cerebellum’s internal prediction is updated, and *β* determines the extent to which the motor control parameter is updated. In the main simulations, the initial values for these parameters were set to *α* = 0.002 and *β* = 0.02, respectively. We ran 12 additional optimizations with different learning and adaptation rate combinations to explicitly investigate the interplay between the two parameters. In these, we fixed the adaptation rate to either a low value (*β* = 0.01) or a high value (*β* = 0.04), while simultaneously setting the initial guess for the learning rate to either a low value (*α* = 0.01) or a high value (*α* = 0.05). The reflex parameters were still optimized. We conducted three parallel optimizations for each of these four combinations with a maximum 80 s simulation length.

### 2.6 Evaluation

We test the effectiveness of our cerebellum model by comparing the SLA between the reflex-only model and the model with the cerebellum. Therefore, for every simulation, we computed the SLA per stride as the difference between the fast and slow step lengths divided by their sum [7]. The parameters were averaged on a stride-by-stride basis across all respective parallel simulations. The number of strides varied across simulations; therefore, if one simulation had more strides than others, only its final stride was excluded to ensure that averaging across simulations on a stride-by-stride basis always included data from at least two simulations. For the simulations with the cerebellum, we additionally decomposed the resulting movements into spatial and temporal components, as proposed by Malone and Bastian [31], to better understand the effect of adapting only the timing of liftoff. The spatial component, defined as the difference between the centers of oscillation of the fast and slow legs, was calculated stride-by-stride by subtracting the center of oscillation of the fast leg from that of the slow leg. The center of oscillation for each leg was determined as the mean of the limb angle at heel strike and toe-off. In our case, the limb angle was computed as the angle between a vertical line and the vector from the pelvis COM to the calcaneus COM. The temporal component, representing the phasing between the legs, was computed for each stride as the lag time at the peak cross-correlation of the fast and slow limb angle trajectories, normalized to the stride duration, with the slow leg serving as the reference. For details about the computation, we refer to [31, 76]. To assess the effect of SLA adaptation on the kinematics, the slow and fast legs’ hip, knee, and ankle joint kinematics of the solutions with cerebellum were normalized to the percentage of the stride and compared between the beginning and end of the simulations, corresponding to early adaptation (EA) and late adaptation (LA). Finally, to evaluate the similarity of our simulations to experimental gait, we calculated the normalized cross-correlation coefficient (NCC) [77], which assesses waveform similarity, and the root-mean-squared error (RMSE), which quantifies the absolute deviation, between the average EA and LA gait cycles from the simulations and our participant’s kinematic data. We also analyze the peak muscle activations of each muscle during EA and LA for both legs. The code and results of this study are publicly available at https://simtk.org/projects/splitbelt, allowing others to replicate our work and build on it.

### 2.7 Statistical analysis

Statistical analyses were performed using the average of the first three strides, corresponding to EA, and the last three steps, corresponding to LA. To assess the effect of the cerebellum model, we conducted a mixed ANOVA with SLA as the dependent variable, time (EA, LA) as the within-subject factor, and simulation type (cerebellum / no cerebellum) as the between-subject factor, testing the hypothesis that adaptation during the split-belt period occurs only in the simulations with the cerebellum. Where significant effects were found, pairwise post-hoc comparisons were performed. To quantify the time course of the adaptations, we fit exponential decay functions *y* = *a − b · e^−s/c^* [7] to the left and right step length, where *s* is the stride number, *c* quantifies the adaptation rate and *|b|* the amount of adaptation. Since we hypothesized that the fast step length would adapt more than the slow step length (Figure 1), we conducted a one-sided t-test comparing *|b|* between both legs. For the spatial and temporal adaptation, we used paired t-tests to compare EA and LA center-of-oscillation and limb phasing values, respectively. For all tests, we set the significance level to 0.05. We applied Bonferroni correction to correct the p-values where multiple comparisons were performed.

## 3 Results

The model was able to maintain a stable walking pattern for all 80 s simulations and showed adaptation patterns when extended with the cerebellum. Videos of a representative 2:1 split-belt walking simulation, both with and without the cerebellum model, are provided in S2 Video and S3 Video, respectively. Within the manuscript, we concentrate on the predicted gait patterns as the most relevant outcome of predictive simulations. Internal cerebellar parameters and variables are provided in S5 File.

### 3.1 SLA adaptation in models with and without cerebellum

Simulations with the cerebellum model demonstrated a gradual adaptation of SLA, starting at -0.17 at the beginning of the adaptation and improving to -0.03 by the end (Figure 4). In contrast, when the cerebellum was not included, the step lengths remained asymmetric at around -0.15 throughout the entire simulation period (Figure 5). ANOVA revealed significant main effects of time (*F*_1,18_ = 78.22, *p <* 0.001) and group (*F*_1,18_ = 11.74, *p* = 0.003), as well as a significant time × group interaction (*F*_1,18_ = 100.13, *p <* 0.001). Post-hoc comparisons confirmed a significant decrease in SLA from start to end across time (*t*_19_ = 3.55*, p* = 0.009) and a significant difference between cerebellum and no-cerebellum simulations (*t*_18_ = 3.43*, p* = 0.012). At the end of adaptation, the SLA differed significantly between groups (*t*_18_ = 8.12*, p <* 0.001), while no difference was observed at the start (*t*_18_ = *−*0.88*, p* = 1.0). These findings show that SLA adaptations occured with the cerebellum model, but not with the spinal controller alone. Our results are consistent with experimental findings from studies involving healthy participants and patients with cerebellar damage [15], where patients exhibit an asymmetric gait pattern at the onset of the split-belt perturbation but do not show adaptations over time. The adaptation pattern is similar to what is reported in [30, 78], with a high SLA change rate during the first strides that decreases over time.

**Fig. 4.**
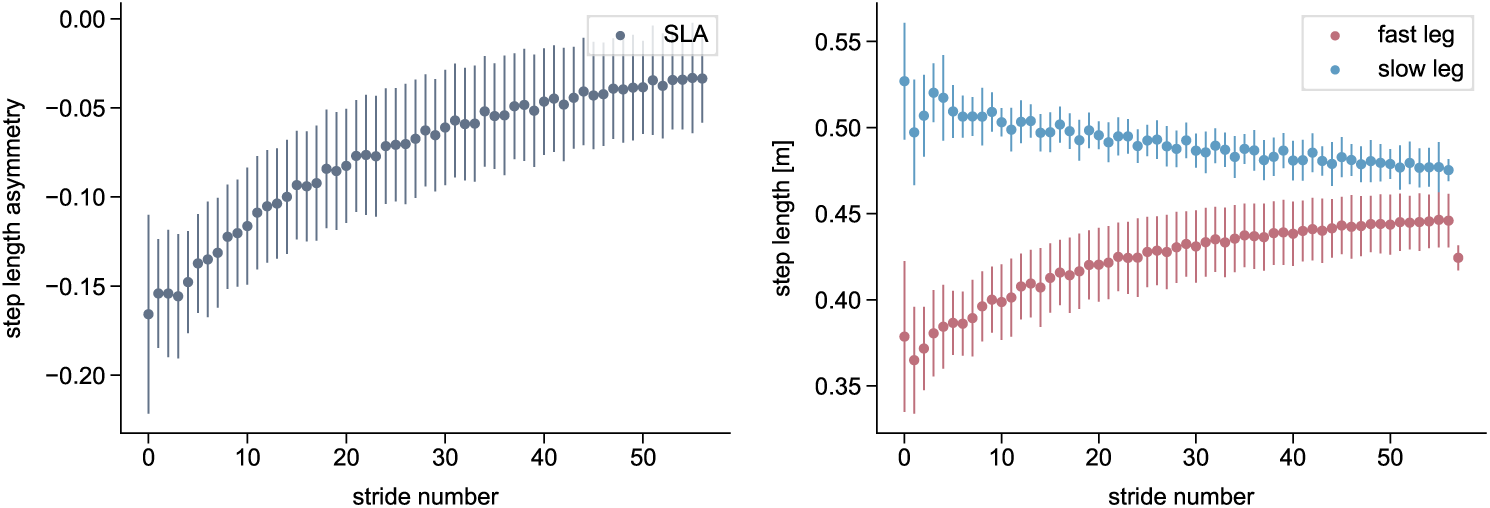
Simulated step length asymmetry and step lengths of the reflex model with the cerebellum during split-belt treadmill walking at a 2:1 speed ratio. All values are averaged over the 80 s simulations, with the bars representing the standard deviation.

**Fig. 5.**
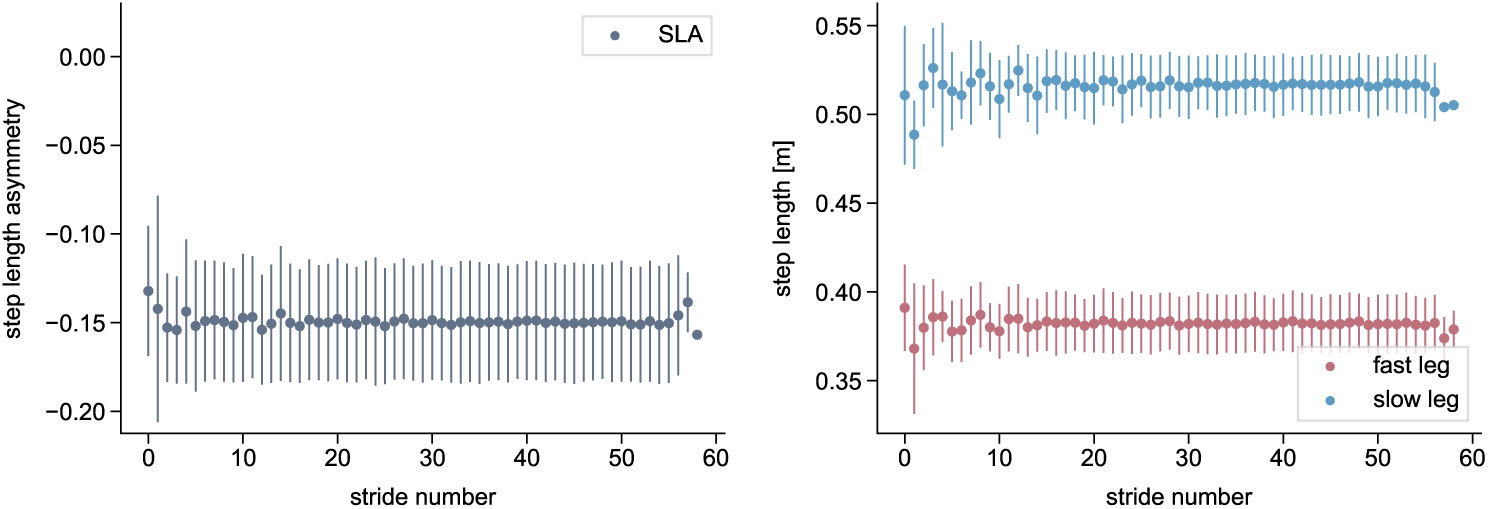
Simulated step length asymmetry and step lengths of the reflex model without the cerebellum (spinal gait controller only) during split-belt treadmill walking at a 2:1 speed ratio. All values are averaged over the 80 s simulations, with the bars representing the standard deviation.

As hypothesized, the adaptation magnitude *|b|* was significantly greater for the fast step length than for the slow step length (*t*_9_ = 3.17, *p* = 0.006). Correspondingly, step length increased for the fast leg from 0.37 m to 0.45 m and decreased slightly for the slow leg from 0.51 m to 0.48 m, consistent with previous reports indicating that the fast leg contributes more strongly to adaptation [7, 79, 80].

### 3.2 Spatial and temporal adapatations

The center of oscillation difference and limb phasing showed significant changes between EA and LA (*t*_9_ = 3.19*, p* = 0.022; *t*_9_ = 5.88*, p <* 0.001, respectively) (Figure 6). Thus, both spatial and temporal components contributed to the SLA adaptation caused by isolated modulation of the swing initiation threshold. This supports the idea of a hierarchical relationship between the temporal and spatial domains, with temporal adjustments affecting spatial parameters [32]. Our simulated limb phasing qualitatively aligns with experimental findings, in which the phasing was out of phase at the start of the adaptation period, with the fast leg leading the slow leg, before rapidly returning toward baseline [31]. However, the simulated center of oscillation difference decreased rather linearly, in contrast to the exponential decay typically seen in humans [31].

**Fig. 6.**
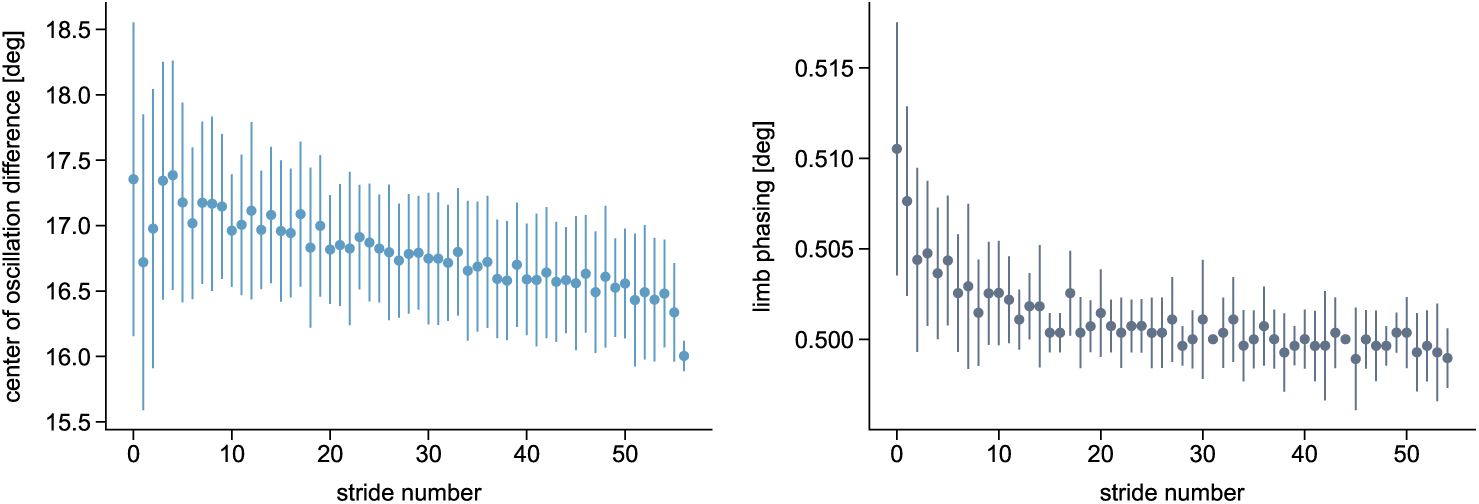
Spatial and temporal adaptation components. The spatial (left plot) and temporal (right plot) components are defined as the center of oscillation difference between the limbs and the interlimb phase, respectively, with the latter computed as the lag time at the peak correlation of each cycle [31]. The baseline values for symmetric walking are 0*^◦^* for the center of oscillation difference and 0.5 for the limb phasing. The values are averaged over the ten 80 s simulations with the cerebellum, with the bars representing the standard deviation.

### 3.3 Validity and adaptation of kinematics and muscle activations

The shape of the simulated hip and knee joint angle trajectories during EA and LA (Figure 7) matched the representative experimental data well, with all NCC values above 0.77, and all values for the fast leg even above 0.96 (Table 1 from S6 File). The slightly lower NCC values in the slow limb were likely due to the straight knee during stance and the increased hip flexion during swing. While the slow ankle also showed good agreement (NCC *>* 0.7), only the fast ankle waveform deviated strongly from the experimental pattern (NCC *<* 0.13). The RMSE values ranged between 5.3°(slow ankle LA) and 14.9°(slow hip LA), indicating an offset in certain absolute joint angle values from the experimental data (S6 File). In humans, the shift from symmetric walking to split-belt walking induces asymmetries between the kinematics of both limbs, particularly at the ankle joint [81], whereas the kinematics of each joint remain largely unchanged throughout the 2:1 split-belt perturbation period [12]. This is well reflected by our model, as all kinematic trajectories differed between the fast and slow limb, but showed little changes over time (most range of motion differences between EA and LA *<* 1.5°, only for slow knee 3.3°). The timing of peak hip and knee flexion and ankle plantarflexion during toe-off qualitatively matched experimental observations, including the slight phase shifts over time: peaks occurred slightly earlier in the slow leg and later in the fast leg during LA, consistent with our participant’s data and previous reports [7].

**Fig. 7.**
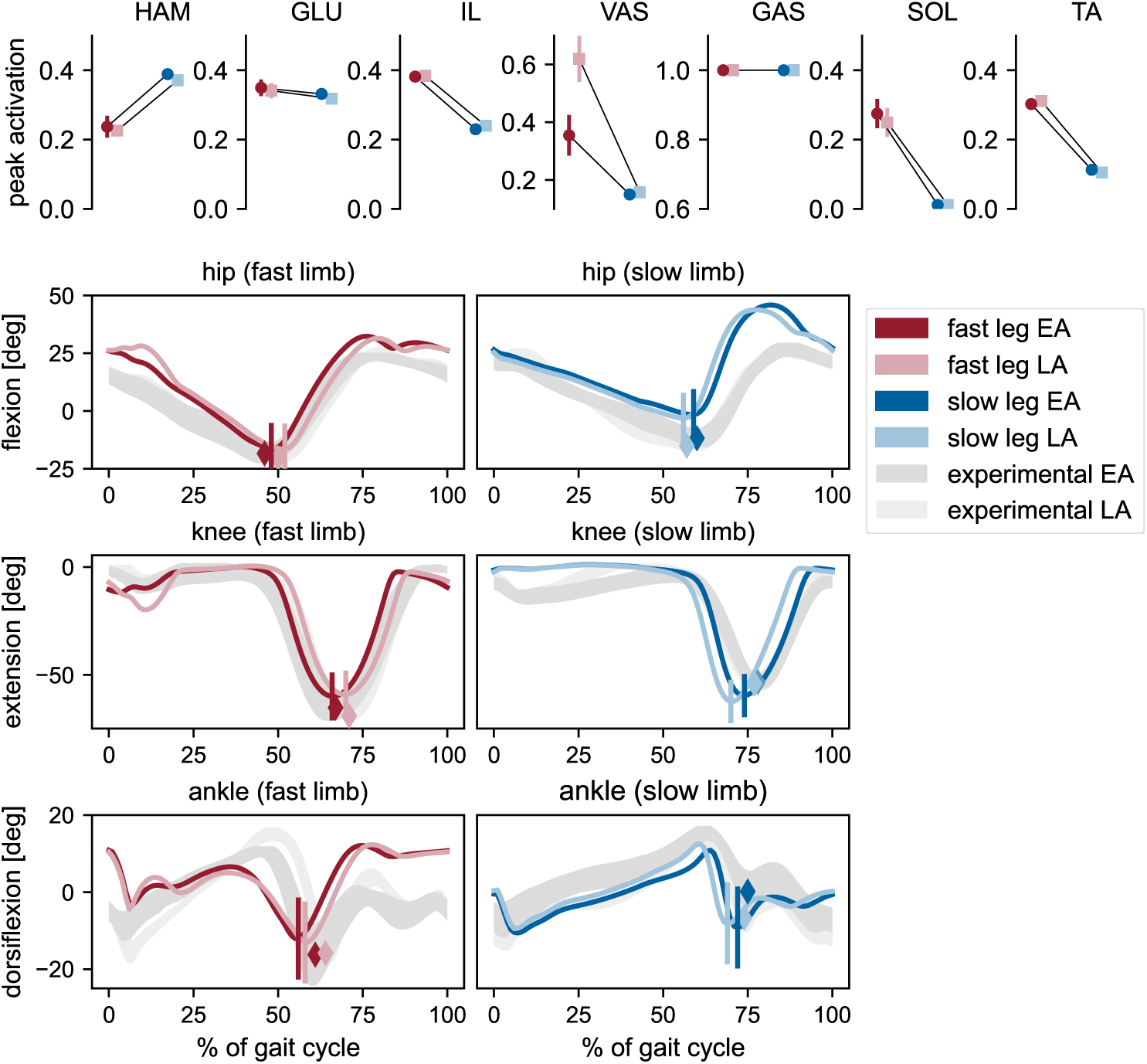
Peak muscle activations and joint kinematics of simulations with cerebellum. Upper row: Peak activations of each muscle for the fast and slow limb during early adaptation (EA) and late adaptation (LA), computed from the first and last five strides, respectively. Dots and error bars represent the mean and standard deviation across ten 80 s simulations. Lower rows: Joint angle trajectories of the fast and slow hip, knee, and ankle joints during EA and LA. Colored lines show the average of the 10 simulations. Shaded areas represent experimental values (mean *±* standard deviation of the first and last 20 strides from one representative participant). Vertical lines indicate the time and value of peak hip extension, peak knee flexion, and peak ankle extension during pushoff in the simulations. Colored diamonds mark the corresponding peaks in the experimental data.

Most peak muscle activations showed differences between the slow and fast limbs, consistent with literature reporting distinct adjustments to the split-belt condition in each leg [35, 36]. Only the gastrocnemius reached its maximal activation in both limbs, with patterns showing saturation at this level (S6 File), which hints at an underestimated maximum muscle force. Similar to experimental findings, lower limb muscle activity was generally higher in the fast leg compared to the slow leg [35, 36, 82], although it has also been suggested that for the upper limb, vastus activity in the slow leg may exceed that of the fast leg [35]. Most muscle activation patterns were similar between early and late adaptation, as expected given the absence of reflex gain adaptation, though the fast vasti showed a noticeable increase in peak activation in LA compared to EA. This likely reflects the increased knee extension moment due to longer step lengths, resulting in greater vasti force, which in turn drives stronger reflex activation despite unchanged reflex gain settings. Increased peak vasti activations for increasing step lengths have also been observed elsewhere [61, 83].

### 3.4 Interplay between learning rate and adaptation rate

The adaptation pattern in our simulations was influenced by both the adaptation and learning rates (Figure 8). For a given adaptation rate, the learning rate determined the SLA level to which the model converged in LA, with lower learning rates resulting in a more positive SLA. The combination of high adaptation and low learning rate even reached a positive SLA level. Conversely, for a fixed learning rate, the adaptation rate affected the speed of adaptation, where higher adaptation rates resulted in a steeper initial increase of SLA during the first strides. In our model, the final asymmetry level is reached when the cerebellar error signal becomes zero, signaling that the model has established a new internal equilibrium or “new normal.” How fast this occurs depends on the specific combination of adaptation and learning rates. The high learning rate condition led to rapid convergence for both adaptation rates. Notably, even the low adaptation rate combined with a high learning rate achieved fast convergence. In contrast, the combination of low adaptation and low learning rates resulted in noticeably slower convergence between prediction and observation. In this case, the cerebellar error signal—and thus the adaptation—did not fully resolve within the 80 strides simulated (see Fig 3 from S5 File). Jacobsen and Ferris [84] have shown that fast and slow adapters exist and that these groups are related to differences in brain activity. The adaptation curves that they found experimentally for fast- and slow-adapting individuals qualitatively match our results for high and low adaptation rates with a low learning rate, in the sense that fast learners show a more exponential adaptation curve and reach a steady-state level of SLA with fewer steps than slow learners.

**Fig. 8.**
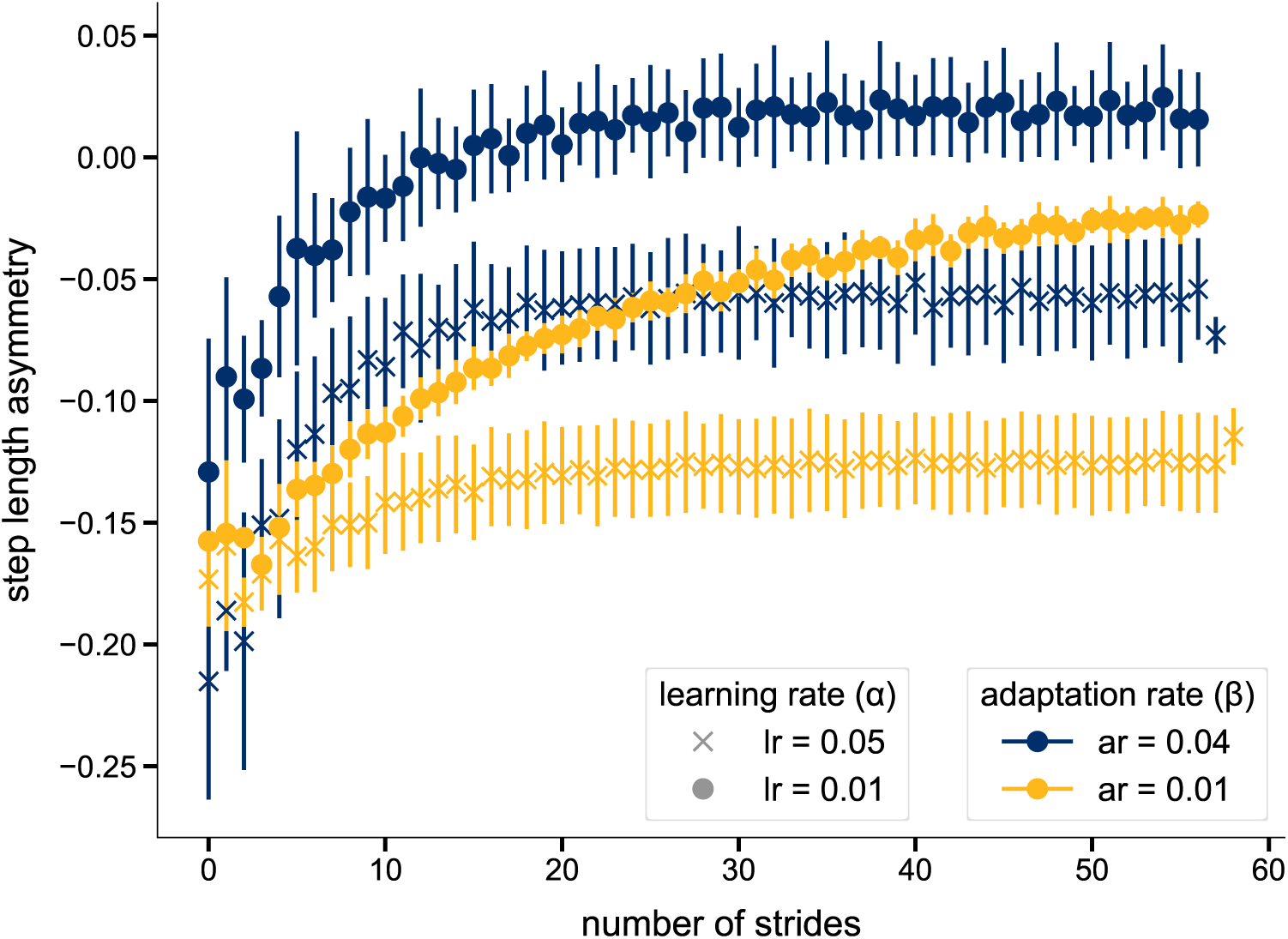
Step length asymmetries for different adaptation and learning rates. The values are averaged over three 80 s simulations for each combination of adaptation and learning rates. The bars represent the standard deviation.

## 4 Discussion

We proposed a biologically inspired model of the cerebellum that learned from prediction errors and could be integrated with a commonly used spinal reflex model as a supraspinal control layer that autonomously generated feedforward adaptations. When combined, the model exhibited SLA adaptations consistent with human split-belt walking behavior, while the spinal reflex-only model was able to walk on the split-belt treadmill but showed no adaptations. The extended model could adapt at different exponential rates and converge to different SLA levels, including positive values, reflecting the variability observed across individuals in experimental data.

Our work complements the split-belt simulation study by Refy et al. [53], who showed that reflex gain changes during split-belt walking could cause SLA adaptations. Here, we found that SLA adaptation could also be driven by the modulation of timing alone, and that, in simulation, reflex changes were not necessarily required for step length adaptations during split-belt walking. Only the initial asymmetries at the beginning of the split-belt period did require adjustments in reflex parameters relative to symmetric walking to stabilize gait, which was the same for Refy. Even though we prohibited feedforward reflex modulations in our experiments, we do not want to contradict that reflexes change during split-belt exposure, especially since the gastrocnemius H-reflex, and also muscle activations, which are a consequence of the spinal reflexes, have also been shown to adapt in experimental studies [36, 54]. Instead, we used the simulations to focus on a single mechanism and isolate its contribution to SLA adaptation. From the work, we cannot conclude whether one mechanism is correct over the other, and split-belt adaptations may be the result of a combination of both reflex changes and timing adaptations.

Neuromusculoskeletal simulations are a powerful tool for determining the individual contribution of interdependent and co-occurring processes. With our simulations, we were able to investigate the role of gait timing adaptation, specifically the start of swing initiation, while blocking reflex changes during the split-belt exposure. Decomposing the simulations into spatial and temporal components showed that restricting adaptation to timing changes resulted in exponential changes in the temporal domain, but only linear changes in the spatial domain. In human experiments, both domains typically adapt exponentially, albeit at different rates. The fact that the spatial domain changed at all, despite only altering timing, may support the proposed hierarchical relationship between temporal and spatial control [32]. The results further suggest that additional control mechanisms, such as reflex gain changes, may be required to reproduce the full spatial adaptation observed in humans.

We model cerebellar adaptations as an error-driven process. While this is supported by many studies [7, 26, 29, 85], locomotor adaptation has also been attributed to energy optimization [6, 86]. During split-belt walking, the gradual SLA reduction is accompanied by a reduction in mechanical work performed by the legs [87] and a reduction in whole-body metabolic cost, likely driven by a reduction of muscle activity [27]. In addition, humans can learn to accept the assistance from the split-belt treadmill and converge to positive SLA values that are energetically advantageous [13, 14], even though they do not reach the same extensive levels that are predicted as optimal [40]. While it is indubitably true that muscle activations and metabolic cost, which increase at the onset of the split-belt perturbation, decrease throughout the adaptation period, it remains unclear if the cerebellum is involved in these changes. Seethapathi et al. [6] have adopted this theory of energy minimization driving locomotor adaptations and successfully predicted human-like SLA adaptations beyond symmetry in a biped walker. However, because step length was a direct control variable of the model, and a nominal step length value was both an input to the controller and updated during adaptation, this finding primarily reaffirms that positive SLAs are energetically beneficial. In our work, we assume that energy minimization is not cerebellar-driven, but rather a separate process within the CNS that contributes to adaptation. This perspective is supported by studies reporting no direct relationship between spatiotemporal adaptations and energy reduction [10, 78]. Furthermore, in a study in which the participants’ energetic landscape during tied-belt walking was altered, the participants did not automatically alter their gait to reach the more economical walking pattern [66], which should have happened, though, if energy was considered the error function of the cerebellum. We have incorporated an effort term into the objective function to reflect the human preference for energy-efficient gait patterns, but this does not alter the fundamentally error-driven nature of cerebellar adaptation in our model.

The error in our cerebellum model was computed from the difference between a predicted and observed SVA. SVA has recently been proposed as the potential target function on split-belt adaptations [29]. It overcomes the limitation that a minimization of SLA, which has often been considered the error signal, would never yield positive SLAs as experimentally observed [6, 88] and that symmetric step lengths are not energetically optimal [86]. Indeed, our model was able to converge to a positive SLA level when the adaptation rate was high, but it would also converge to slightly negative SLA levels in other simulations. In contrast, when we used SLA as the error metric, the simulations never reached symmetry or positive SLA values. This shows that the target function proposed by Ishida et al., which was derived from the goal-equivalent manifold and is related to decreasing kinetic energy [29], works, also in simulation. However, other potential error metrics, such as the minimization of double support time, could still be tested.

Closely related to the definition of the error signal is the structure of the prediction and error computation in our cerebellum model. We used one interlimb error metric, from which the updates for both limbs were derived in a gradient-based manner. This is different from Fujiki et al. [51], who computed a distinct intralimb error for each limb and directly used the error in calculating the duty factor of the respective robotic leg. Malone et al. [26] even proposed different error metrics for spatial and temporal adaptations, based on the long-held assumption that these two domains are controlled independently, even though recent studies have challenged this view [32].

In our work, we used the same adaptation rate for both limbs. Even though this captured the experimental finding that the fast leg contributes more to SLA adaptation than the slow leg, separate control circuits may exist for the left and right legs. In mice, temporal adaptation was only impaired when chemogenetic manipulation was applied to the cerebellar hemisphere on the same side as the leg undergoing the speed change, but not when the opposite side was targeted [85]. The presence of two neural circuits for left and right, rather than a single one, might also explain why the predicted swing initiation threshold of the slow leg in our simulations qualitatively showed more pronounced changes than suggested by the experimental data, and should be tested in follow-up studies.

The biological interpretation and validity of the learning and adaptation rate in our computational model remain open to discussion. Depending on the rates, our model predicted differences in the SLA adaptation curves, which have also been observed experimentally. While Jacobsen et al. [84] attributed such differences to fast and slow learners, another study suggests that adaptation rates, and consequently the differences in the exponential decrease of error, can be influenced by prior experience [89]. Applied to this study, this would mean that the adaptation rate in our model could either remain fixed for a given individual or vary with context. Another question that arises and should further be investigated is whether the update rate of the cerebellar prediction and the update rate of motor commands are actually separable in humans, as proposed in our work, and can thus be modulated or trained independently, or if they are inherently linked. Ultimately, our cerebellum model is a computational abstraction of the much more complex biological structure, meaning that the concept of a single learning and adaptation rate, while effective in simulation, will likely not correspond exactly to reality.

While the waveforms of the predicted joint kinematics and their differences between EA and LA were mostly similar to experimental data, the fast ankle joint trajectory, as well as the RMSEs, displayed some deviations. There are multiple potential reasons for this, such as the design of the spinal controller, which may not be ideally suited for split-belt treadmill walking, or the objective function that we used to optimize the spinal parameters. While the objective function incorporated different aspects relevant for human gait, we cannot be certain that these terms are equally relevant for split-belt walking. It has been shown, for example, that maintaining stability against the continuous perturbation induced by the two belt speeds is particularly relevant in split-belt walking [90], specifically at the beginning of the exposure [78]. However, those kinematic deviations do not affect our primary finding, which was the ability of our cerebellum model to predict locomotor adaptation in a reflex model. Notably, there is also other modeling work that successfully predicted gait changes, for instance, in patients with neural impairments [48, 70] or due to sensory modulation [61], despite observed mismatches to experimental joint-level kinematics. Future efforts should, however, be dedicated to improving the spinal control part of the model to further enhance the realism of the predicted adaptations.

Our study has several limitations. One limitation is that we had to manually read-just certain reflex gains when determining the initial guess for the final simulations, as the parameter regions leading to experimentally observed asymmetry levels were not reached automatically. However, it is not uncommon in predictive simulations that the initial guess is tuned by hand, specifically for novel tasks for which no previous solution is available [50, 56, 91]. Other studies do not report the initial guess at all, which suggests that manual tuning may also have been used in those cases [92, 93]. Nevertheless, more elaborate methods might be applied to find a good initial guess, such as an imitation objective that finds the initial guess parameters by minimizing the difference between predicted and experimental states. In addition, we started with an initial guess that already incorporated the reactive changes that occur immediately when the belt speeds are made asymmetric. Thus, our model does not capture the rapid, reactive responses at the transitions between baseline and early adaptation or between late adaptation and washout, nor how these responses are shaped by learning or recalibration of an internal model. Future work could extend the study and also include these transitions from and to symmetric belt speeds, as was done for the biped model in [6], and also include after-effects upon returning toward tied belt speeds. Rather than directly updating the prediction generated by the internal model, future studies should investigate what information is stored and how the prediction is derived from the internal model. Another limitation is that our model currently does not explain the experimentally observed changes in muscle activations or center-of-oscillation differences. Since we only adapted the timing of swing initiation, we can conclude that these changes are likely not driven by timing alone. However, we can only speculate about the control structures responsible for them, such as modifications in reflex parameters.

Our study focused on split-belt walking, because it is the most common paradigm to investigate locomotor adaptations as previously stated by [1, 6]. Moreover, the paradigm has clinical relevance in the field of neurorehabilitation [8, 94–97], which makes it particularly important to gain a detailed understanding of the individual components and mechanisms underlying locomotor adaptation. By choosing this specific task for our study we could show that the reflex-based model alone, without supraspinal control like our cerebellum model, is not sufficient to capture human behaviour, which would not have been obvious if we limit the study to general tasks like normal walking. Nevertheless, future work should consider additional locomotor tasks and incorporate additional or more detailed supraspinal circuits. This would contribute to the development of a neuromusculoskeletal model capable of generalizing and explaining various types of behaviors.

Predictive simulations always work with idealized computational models, which are based on assumptions and might not capture all control structures of a human. Our neuromusculoskeletal model was a reflex model that is frequently used in literature and has demonstrated robustness against other types of perturbations. However, different models could be better suited for the investigation of specific aspects of motor adaptation in split-belt walking. For example, Malone et al. [26] have suggested that adaptations in the spatial domain are a consequence of modulating foot placement, but active foot placement is not possible with our spinal reflex model. Song and Geyer [50] have proposed a neuromusculoskeletal model with an additional supraspinal control structure that adjusts the desired foot placements and could serve as an alternative to our approach of adapting the start of the swing initiation. Furthermore, there are indications that CPGs might contribute to the spinal control of locomotion [98]. CPGs are neural circuits that generate rhythmic motor patterns without the need for sensory feedback. These circuits have been integrated into neuromusculoskeletal models both on top [99] or instead of [47] reflex-based neural circuits. Such models could help explore whether the phase-resetting approach proposed by Fujiki et al. [51], which was developed for a torque-controlled robot to replicate gait adaptations, could be generalized to a neuromusculoskeletal model. However, our results demonstrate that even a basic reflex model is sufficient for walking on a split-belt treadmill and for effectively capturing salient adaptation features.

Our work incorporates a more physiologically detailed model and control structure than previous split-belt simulation studies that relied on simplified models with low-dimensional or torque-based control strategies [6, 51, 52]. Although such models can be used to explain general concepts of locomotion, their control architectures and dynamic model parameters are not transferable to the human neuromuscular system, which comprises multiple segments and an overactuated system of muscles. Consequently, they offer limited insight into how motor adaptation might be physiologically realized. Nevertheless, the model by [6] could generalize to a wider range of tasks and conditions, which still remains an open challenge for our neuromusculoskeletal model.

To our knowledge, this is the first study in which a computational model of the cerebellum autonomously drives feedforward gait adaptations in a neuromusculoskeletal model in response to the continuous perturbation imposed by a split-belt treadmill. By interacting directly with spinal reflex circuitry, the model enables the continuous interplay between supraspinal and spinal control circuits and the musculoskeletal system. Beyond step length asymmetry, we report adaptations across multiple domains, including spatial and temporal contributions as well as kinematics, providing a more comprehensive picture of simulated locomotor adaptation than previous studies [6, 53]. The framework enables disentangling individual mechanisms like, in our case, isolating timing-related parameter changes while blocking reflex adaptation, thereby offering mechanistic insights that are difficult to obtain experimentally. While such computational approaches inherently involve simplifications and abstractions, and results should be interpreted with caution, the framework can be readily extended to test additional hypotheses and thereby support the broader goal of advancing mechanistic understanding and predictive modeling for locomotor rehabilitation.

## 5 Conclusion

In conclusion, we proposed a computational model of the cerebellum, which success-fully generated SLA adaptations in a reflex model during split-belt walking. We found that reflex changes are not strictly required for SLA adaptations, and that changes in the timing of swing initiation are sufficient to explain experimentally observed temporal adaptations, but not spatial adaptations. The model is a step towards integrating supraspinal learning mechanisms with neuromusculoskeletal simulations to investigate the neural control of locomotor adaptation from a simulation perspective.

## Supplementary information

- S1 File *pdf.* Details on the experimental data and parameter computation.
- S2 Video *mp4.* Video of one representative split-belt walking simulations at 2:1 belt speed ratio with adaptations driven by the cerebellum model.
- S3 Video *mp4.* Video of one representative split-belt walking simulations at 2:1 belt speed ratio without adaptations driven by the cerebellum model.
- S4 File *pdf.* Details on the spinal control laws and controller parameters.
- S5 File *pdf.* Additional results on the optimized cerebellar parameters and internal model variables relevant to the simulations presented in the manuscript.
- S6 File *pdf.* Additional kinematic results and muscle activation patterns.

## Abbreviations

CMA-ES: Covariance Matrix Adaptation Evolution Strategy
COM: Center of mass
CNS: Central nervous system
EA: Early adaptation
ES: Early stance
GAS: Gastrocnemius
GLU: Gluteus maximus
HAM: Hamstrings
HS: Heel strike
IL: Iliopsoas
LA: Late adaptation
LP: Landing preparation
MS: Mid-stance
NCC: Normalized cross-correlation
PD: Proportional–Derivative
PS: Pre-swing
RMSE: Root-mean-squared error
SLA: Step length asymmetry
SOL: Soleus
SV: Step velocity
SVA: Step velocity asymmetry
SW: Swing
TA: Tibialis anterior
VAS: Vasti

## Declarations

### Ethics approval and consent to participate

The study was approved by the ethical committee of the Friedrich-Alexander-Universität Erlangen-Nürnberg (Re.-No. 22-437-B). The participants gave informed consent prior to participation and was informed about the regulations of participation. All methods were carried out in accordance with relevant guidelines and regulations.

### Consent for publication

Written informed consent for publication of anonymized data was obtained from the participant.

### Availability of data and materials

The datasets generated and analysed during the current study are available in the SimTK repository https://simtk.org/projects/splitbelt. The repository contains all data, simulation results and code, and analysis scripts necessary to reproduce the findings reported in this manuscript.

### Authors’ contributions

SF conceptualized the work. SF, CO and AD developed the methodology and models. SF conducted the experiments and analyzed the results. CO, BME and ADK supervised and mentored the research. SF wrote the paper. All authors thoroughly reviewed, edited and approved the manuscript.

## Acknowledgments

We thank Thomas Geijtenbeeck for his support with SCONE and Hyfydy.

