## Supplementary material for "A model of the cerebellum generates gait adaptations in a reflex-based neuromusculoskeletal model during split-belt walking": S1_file

### S1 - Details on experimental data

#### Data collection

We recorded motion capture data from one representative participant while walking on a split-belt treadmill. The participant gave informed consent prior to participation and was informed about the regulations of participation. The study was approved by the ethical committee of the Friedrich-Alexander-Universität Erlangen-Nürnberg (Re.-No. 22-437-B). All methods were carried out in accordance with relevant guidelines and regulations.

We used the same study protocol as Buurke [1] or Hoogkamer [2]. The participant walked on a treadmill with both belts running at 1 m/s for five minutes. During the subsequent 10 min intervention period, the speed of the right belt (since the right leg was the participant's dominant leg) was reduced to 0.5 m/s. Eventually, the right belt speed was returned to 1 m/s, and the subject walked again for 5 min. The treadmill was stopped briefly between the three configurations.

Data collection was performed with the GRAIL system (Gait Real-time Analysis Interactive Lab, Motekforce Link), which included an instrumented split-belt treadmill and an optical motion capture system (VICON, Vero, 12 cameras, 100 Hz). We used a full-body marker set to capture the participant's movement.

#### Data analysis

We focused on analyzing the 10 min split-belt period. The force and marker data were filtered using a second-order Butterworth filter with a 10 Hz cutoff frequency. To compute the spatiotemporal parameters during the intervention, heel strike events were detected from the vertical ground reaction forces using a 20 N threshold. Left and right step lengths were computed as the anteroposterior distance between the ankle markers at the respective heel strike [3]. Step length symmetry $s$ was defined as:

$$s = \frac{\mathrm{SL}_{\mathrm{fast}} - \mathrm{SL}_{\mathrm{slow}}}{\mathrm{SL}_{\mathrm{fast}} + \mathrm{SL}_{\mathrm{slow}}}, \tag{1}$$

where $\mathrm{SL}_{\mathrm{fast}}$ and $\mathrm{SL}_{\mathrm{slow}}$ are the step lengths of the left and right legs, respectively [4].

The start of swing initiation was defined as the time point during the stance phase when the anteroposterior distance between the lateral knee marker and the anterior superior iliac spine marker began to decrease. This was taken as the onset of hip flexion and the subsequent limb forward movement.

Experimental joint angles were obtained through inverse kinematics performed in AddBiomechanics [6] using a three-dimensional full-body musculoskeletal model [5].
