## Supplementary material for "A model of the cerebellum generates gait adaptations in a reflex-based neuromusculoskeletal model during split-belt walking": S4_file

### S4 - Spinal controller

#### Spinal control equations

The spinal controller is a reflex controller, originally proposed by Geyer and Herr [1], and has been frequently used in other predictive simulation studies [2–6]. In this controller, muscle excitations $u$ are generated from muscle length and force feedback, a proportional-derivative (PD) controller, and constant feedforward signals, as described by the following equations [2]:

Muscle length feedback:

$$u_L = \max(0, K_L(l(t - t_D) - l_0)) \tag{1}$$

Muscle force feedback:

$$u_F = K_F F(t - t_D) \tag{2}$$

PD balance controller:

$$u_{PD} = K_P(\theta(t - t_D) - \theta_0) + K_V(\dot{\theta}(t - t_D)) \tag{3}$$

Feedforward stimulation:

$$u_C = K_C \tag{4}$$

In the equations above, $l$ and $f$ are the normalized muscle length and force, $\theta$ and $\dot{\theta}$ are the pelvis tilt and velocity, and $t_D$ is a time delay. The controller gains $K_L$, $K_F$, $K_P$, $K_V$ and $K_C$ are free parameters and whose values are determined during the optimization, as are the length offset of the stretch response $l_0$ and the desired pelvis tilt angle $\theta_0$ during swing. Mathematically, for the pelvis tilt, it makes no difference whether the desired angle is prescribed or treated as an optimization variable, since the optimized gain $K_p$ can compensate for any fixed choice of the desired angle.

#### Free controller parameters

For our spinal controller, the above equations, together with the phase-dependent controller depicted in Figure 3 of the main manuscript, lead to the following 31 free spinal control parameters per leg:

| | Stance | | | Swing | |
|---|---|---|---|---|---|
| | ES | MS | PS | SW | LP |
| **HAM** | $K_P$, $K_V$, $K_C$ | | | $K_F$ | |
| **GLU** | $K_P$, $K_V$, $K_C$ | | $K_C$ | $K_F$ | |
| **IL** | $K_P$, $K_V$, $K_C$ | | $K_L$, $L_0$ | $K_L$, $L_0$, $K_P$, $K_V$, $\theta_0$,$K_{L(HAM)}$,$L_{0(HAM)}$ | |
| **VAS** | $K_F$, $K_C$, φ | | | | |
| **GAS** | $K_F$, $K_C$ | | | | |
| **SOL** | $K_F$, $K_C$ | | | | |
| **TA** | $K_L$, $L_0$ $K_{F(SOL)}$ | | | | |

Figure 1: **Spinal controller parameters for one leg** The parameter values are found during the optimization. To prevent knee overextension, the vastus reflex is only active when knee flexion is above a certain threshold $\varphi$, which is also optimized. The same parameters are found for the other leg, resulting in 62 optimization variables.

[5] Veerkamp K, Waterval NFJ, Geijtenbeek T, Carty CP, Lloyd DG, Harlaar J, et al.

Evaluating cost function criteria in predicting healthy gait. Journal of Biomechanics. 2021;123:110530. doi:10.1016/j.jbiomech.2021.110530.

[6] Kiss B, Waterval NF, van der Krogt MM, Brehm MA, Geijtenbeek T, Harlaar J, et al. Minimization of metabolic cost of transport predicts changes in gait mechanics over a range of ankle-foot orthosis stiffnesses in individuals with bilateral plantar flexor weakness. Frontiers in Bioengineering and Biotechnology. 2024;12:1369507. doi:10.3389/fbioe.2024.1369507.
