## Supplementary material for "A model of the cerebellum generates gait adaptations in a reflex-based neuromusculoskeletal model during split-belt walking": S5_file

### S5 - Additional Results

#### Cerebellar predictions and errors - main simulations

Fig 1 shows how internal cerebellar prediction is updated throughout the adaptation. The right plot shows the error which is computed from the cerebellum. It can be seen that the error decreases throughout the adaptation period, but has not converged to zero, which shows that the adaptation is not fully completed yet.

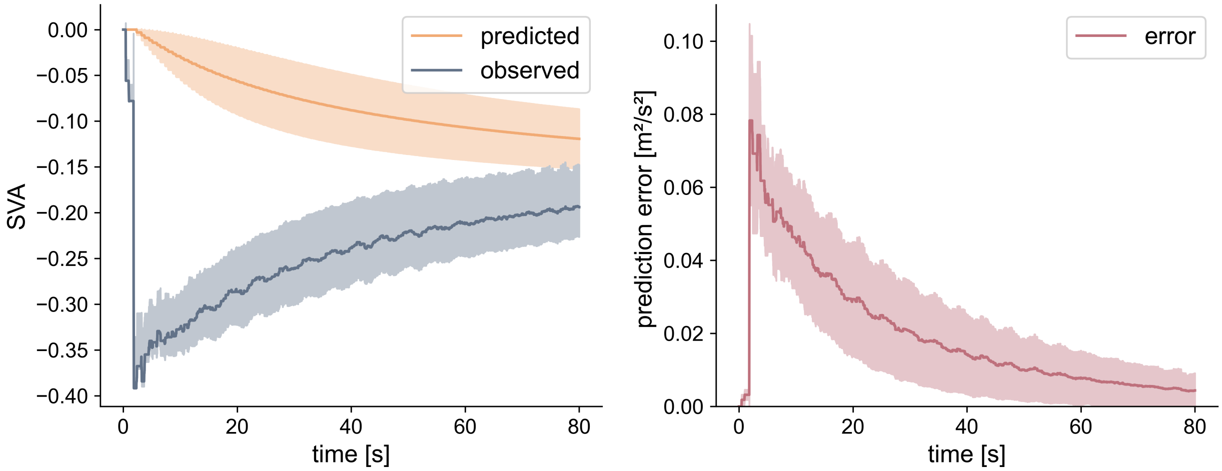


Fig. 1: **Internal model variables and their evolvement throughout adaptation** Left side: predicted step velocity asymmetry (SVA) by the cerebellum and the corresponding observed value over time. Right side: error signal, which is computed based on the prediction and observation. The values are averaged over the 10 main simulations with cerebellum and are plotted as mean ± standard deviation.

#### Optimized cerebellar parameters - main simulations

| Simulation number | Learning rate | Adaptation rate |
|---|---|---|
| 0 | 0.00382 | 0.00841 |
| 1 | 0.009279 | 0.008623 |
| 2 | 0.027530 | 0.023577 |
| 3 | 0.011868 | 0.022930 |
| 4 | 0.002600 | 0.003843 |
| 5 | 0.005035 | 0.008873 |
| 6 | 0.006795 | 0.013375 |
| 7 | 0.004579 | 0.005371 |
| 8 | 0.005357 | 0.002904 |
| 9 | 0.005476 | 0.005154 |

Table 1: Optimized cerebellar learning and adaptation rates for each of the 10 parallel main simulations.

#### Swing initiation thresholds t - main simulations

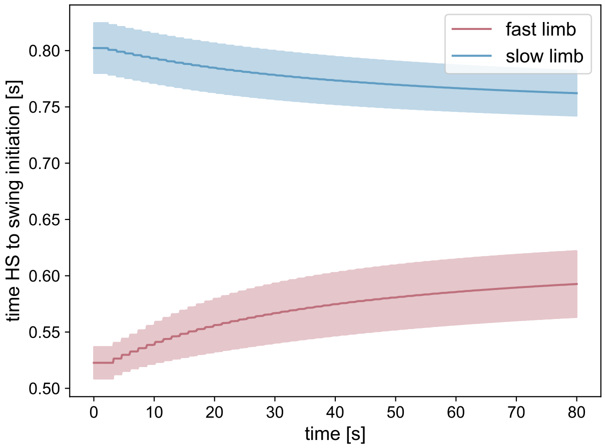


Fig. 2: **Swing initiation thresholds and their evolvement throughout adaptation** The values are averaged over the 10 main simulations with cerebellum and are plotted as mean ± standard deviation.

### Additional experiments - learning and adaptation rate interaction

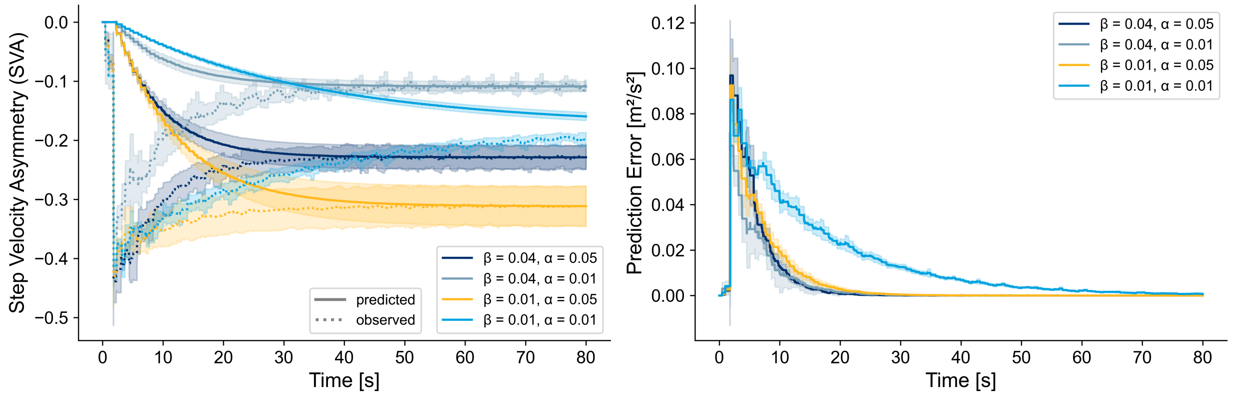


Fig. 3: **Internal model variables and their evolvement throughout adaptation for the different configurations** Left side: predicted step velocity asymmetries (SVA) by the cerebellum and the corresponding observed value over time. Right side: error signal which is computed based on the prediction and observation. The values are averaged over the 3 simulations of each adaptation $\beta$ and learning rate $\alpha$ combination and plotted as mean $\pm$ standard deviation.

| **Adaptation rate $\beta$** | **Learning rate $\alpha$** | **Sim 0** | **Sim 1** | **Sim2** |
|---|---|---|---|---|
| 0.04 | 0.05 | 0.047217 | 0.042891 | 0.047211 |
| | 0.01 | 0.017644 | 0.025421 | 0.032642 |
| 0.01 | 0.05 | 0.050326 | 0.049953 | 0.049785 |
| | 0.01 | 0.009851 | 0.009863 | 0.009888 |

Table 2: **Optimized learning rates for the respective combination of fixed adaptation rate and learning rate initial guess**
