## Supplementary material for "A model of the cerebellum generates gait adaptations in a reflex-based neuromusculoskeletal model during split-belt walking": S6_file

### S6 - Additional Results

#### Kinematics - additional results

| Joint | Leg | Phase | RMSE (°) | NCC |
| --- | --- | --- | --- | --- |
| Hip | Fast | EA | 8.26 | 0.99 |
| | | LA | 7.91 | 0.99 |
| | Slow | EA | 13.57 | 0.86 |
| | | LA | 14.93 | 0.77 |
| Knee | Fast | EA | 6.48 | 0.97 |
| | | LA | 6.92 | 0.96 |
| | Slow | EA | 10.80 | 0.93 |
| | | LA | 13.57 | 0.81 |
| Ankle | Fast | EA | 10.40 | 0.12 |
| | | LA | 11.68 | -0.03 |
| | Slow | EA | 6.17 | 0.75 |
| | | LA | 5.34 | 0.70 |

Table 1: **Root mean squared error (RMSE) and normalized cross correlation (NCC) between the mean simulated data and experimental data from our representative participant.** RMSE compares the absolute values of the mean trajectories, while NCC is a measure of shape similarity.

#### Muscle activations - additional results

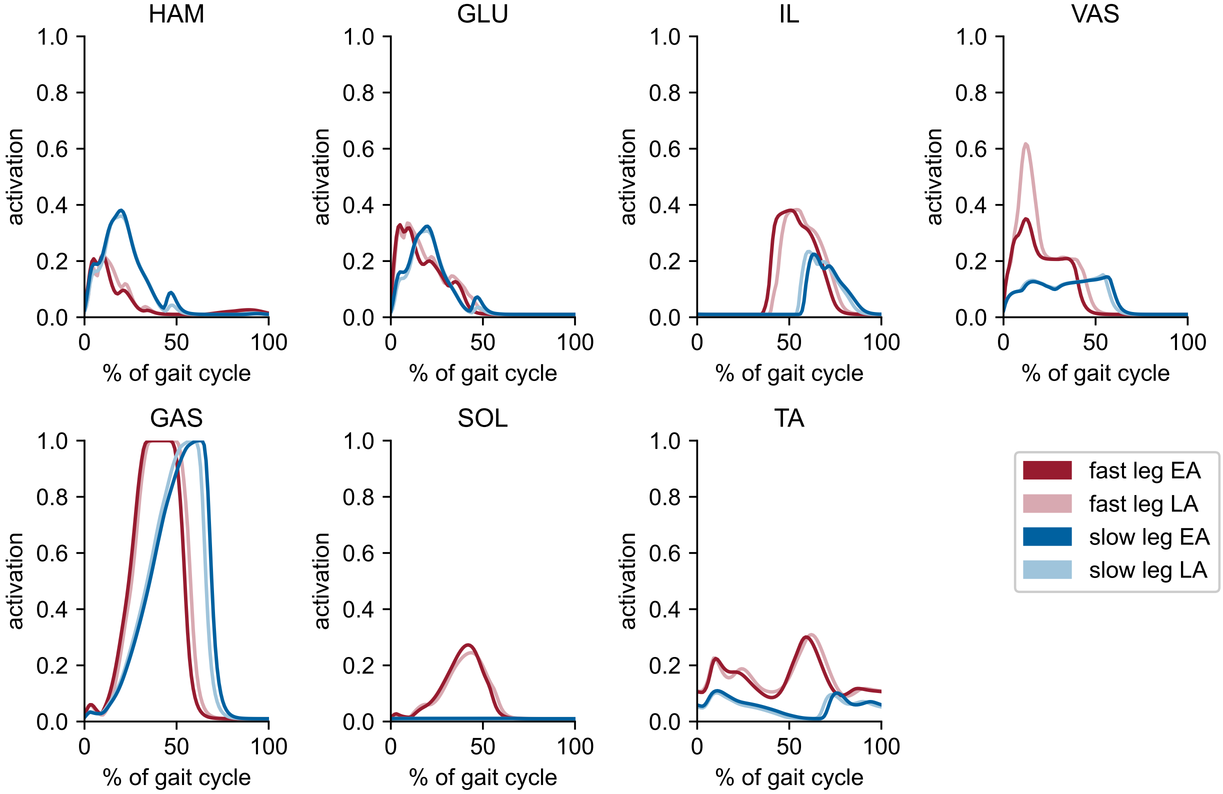


Figure 1: **Muscle activation patterns over time** The trajectories show the average across the 10 simulations with cerebellum.
